# Roles for the RNA polymerase III regulator MAFR-1 in regulating sperm quality in *Caenorhabditis elegans*

**DOI:** 10.1101/2020.04.27.064121

**Authors:** Amy M. Hammerquist, Sean P. Curran

## Abstract

The negative regulator of RNA polymerase (pol) III *mafr-1* has been shown to affect RNA pol III transcript abundance, lipid biosynthesis and storage, progeny output, and lifespan. We deleted *mafr-1* from the *Caenorhabditis elegans* genome and found that animals lacking *mafr-1* replicated many phenotypes from previous RNAi-based studies, but strikingly not the oocyte-related reproductive phenotypes. Utilizing a yeast two-hybrid assay, we discovered several novel interactors of MAFR-1 that are expressed in a sperm- and germline-enriched manner. In support of a role for MAFR-1 in the male germline, we found *mafr-1* null males have smaller spermatids that are less capable in competition for fertilization; a phenotype that was dependent on RNA pol III activity. Restoration of MAFR-1 expression specifically in the germline rescued the spermatid-related phenotypes, suggesting a cell autonomous role for MAFR-1 in nematode male fertility. Based on the high degree of conservation of Maf1 activity across species, our study may inform similar roles for Maf1 and RNA pol III in mammalian male fertility.

## INTRODUCTION

Canonically characterized as a negative regulator of RNA polymerase III, MAF1 was originally discovered and has been extensively studied in *Saccharomyces cerevisiae* (Boguta et al., 1997; Pluta et al., 2001; Upadhya et al., 2002). Since its discovery in *S. cerevisiae*, MAF1 has been identified across diverse eukaryotic clades (Johnson et al.; McLean and Jacobs-Lorena; Pernas et al.; Pluta et al., 2001; Soprano et al.; Upadhya et al., 2002). Perturbation of MAF1 activity leads to, in addition to increased RNA pol III activity, an increase in intracellular lipid abundance (Khanna et al., 2014; Mierzejewska and Chreptowicz, 2016; Palian et al., 2014) and in some instances altered lifespan(Cai and Wei; Shetty et al.). The majority of studies of MAF1 have been conducted in single-cell systems (as *S. cerevisiae*) or cultured mammalian cells (Chen et al., 2018; Johnson et al., 2007; Palian et al., 2014; Rohira et al., 2013), but relatively few studies have probed the function of MAF1 within the context of a complete animal (Bonhoure et al., 2015; Khanna et al., 2014; Pradhan et al., 2017; Rideout et al., 2012).

In *Caenorhabditis elegans*, RNA interference (RNAi) knockdown of MAF1 homolog *mafr-1* results in increased RNA pol III activity, increased intestinal lipid accumulation (as well as increased expression of lipid biogenesis genes), and increased expression of the vitellogenin family of lipid transport proteins (Khanna et al., 2014). As vitellogenesis is the process by which lipids are transferred to developing oocytes, these findings imply a role for MAFR-1 in reproduction, but this has not yet been fully investigated.

Although RNA pol III activity has, to our knowledge, not been directly implicated in fertility, many processes affected by RNA pol III activity do affect reproductive output. *Drosophila* with disrupted ribosome biogenesis display a *Minute* phenotype characterized by delayed development, short and thin bristles, and impaired fertility and viability (Marygold et al., 2007). Furthermore, mice lacking *Zfn384*, a protein whose sub-cellular location was highly correlated with that of RNA pol III in human embryonic stem cells, have fertility defects and impaired spermatogenesis (Alla and Cairns, 2014; Nakamoto et al., 2004). While mature sperm are widely accepted to be transcriptionally and translationally quiescent (Johnson et al., 2011; Kierszenbaum and Tres, 1975), MAFR-1 as a repressor of RNA pol III activity may still be acting in developing germ cells to prevent erroneous RNA pol III transcription.

In all sexually reproducing species, sperm are produced in great excess of oocytes, and must compete with each other to successfully fertilize an oocyte. Generally, larger sperm are able to travel faster than smaller sperm, and therefore have a competitive advantage (Gomendio and Roldan, 1991; LaMunyon and Ward, 1998, 1999). As a hermaphroditic species, *C. elegans* presents a unique situation. The sperm produced by hermaphrodites are capable of fertilizing oocytes but are almost completely outcompeted by male sperm, when mated (LaMunyon and Ward, 1995). In this work, we explore the role of MAFR-1 in maintaining male sperm quality in *C. elegans*, which affects sperm quality metrics such as size and activation capacity, as well as competitive advantage over hermaphrodite sperm.

## RESULTS

### Characterization of a *mafr-1* null mutant

Previous studies of *mafr-1* in *C. elegans* have utilized RNAi-based approaches (Khanna et al., 2014) and a gain-of-function allele of *mafr-1* (Pradhan et al., 2017) leaving the true loss-of-function phenotype unknown. To better examine the biological functions of MAFR-1, we assessed the impact of a true molecular null allele of *mafr-1*, hereafter referred to as *mafr-1* (KO) (**Fig. 1*A***). Surprisingly, a total loss of *mafr-1* did not result in changes in developmental timing (**Fig. 1*B***) or overall organismal health, as *mafr-1* (KO) animals had similar lifespans to wild type (WT) controls (**Fig. 1*C***). Consistent with its canonical role as a negative regulator of RNA pol III activity and previously observed phenotypes for *mafr-1* RNAi (Khanna et al., 2014; Pluta et al., 2001; Upadhya et al., 2002), *mafr-1* (KO) animals showed increased expression of RNA pol III transcripts, including three tRNAs: initiator Methionine, Tryptophan, and Iso-leucine (**Fig. 1*D***). Similarly, *mafr-1* (KO) animals displayed increased intracellular lipid abundance relative to age-matched WT control animals (**Fig. 1*E***), as expected from previous studies (Khanna et al., 2014; Mierzejewska and Chreptowicz, 2016; Palian et al., 2014; Pradhan et al., 2017). Our previous investigations revealed that overexpression of *mafr-1* can influence reproductive output (Khanna et al., 2014). Although the total reproductive output between *mafr-1* (KO) animals and WT controls was unremarkable (**Fig. 1*F***), peak reproductive output appeared delayed in *mafr-1* (KO) animals (**Fig 1*G***). Surprisingly, while the expression of vitellogenins, which deliver lipids from the intestine to the germline to drive reproduction, was previously demonstrated to increase in *mafr-1* RNAi treated animals and diminished by *mafr-1* overexpression (Khanna et al., 2014), vitellogenin expression was not measurably altered in *mafr-1* (KO) animals relative to WT controls at either the transcriptional or protein levels (Fig. S1*A*-*B*). Taken together, these results indicate that *mafr-1* (KO) animals share several of the previously documented *mafr-1*-associated phenotypes, without compromised overall health.

**Figure 1.**
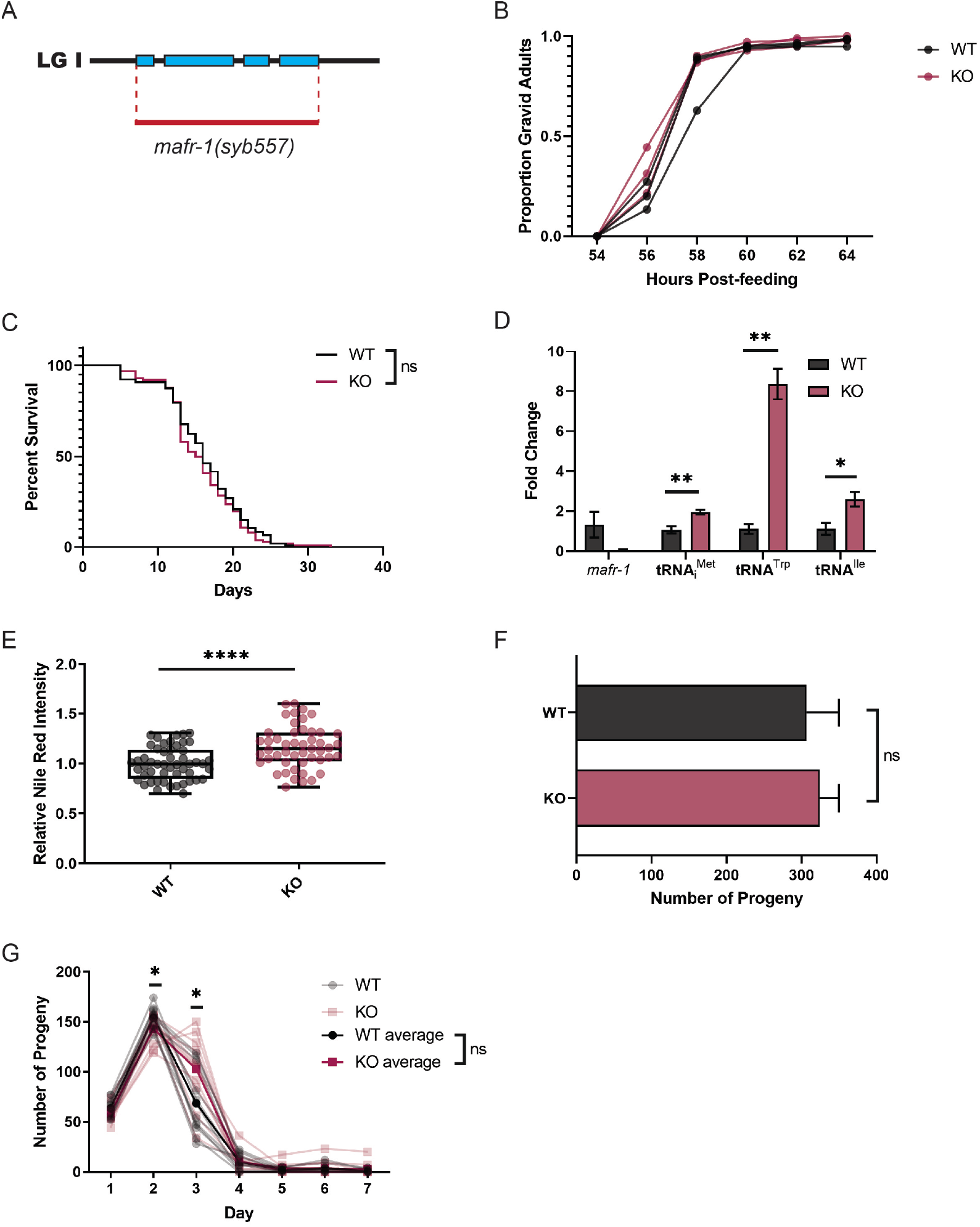
*mafr-1* (KO) produces fertile, viable hermaphrodites. (A) Schematic diagram showing region deleted in *mafr-1(syb557)*, hereafter referred to as *mafr-1* (KO). (B) Developmental time course of WT and *mafr-1* (KO) hermaphrodites. Each line represents one population of >100 animals. (C) Lifespan survival assay of WT and *mafr-1* (KO) hermaphrodites. Individual lines represent average of three 50-worm populations. No significant difference was found by Log-rank (Mantel-Cox) test. (D) Quantitative PCR expression analysis of *mafr-1* and three tRNA RNA pol III transcripts: initiator Methionine, Tryptophan, and Iso-leucine. (E) Lipid abundance as measured by Nile Red staining intensity. (F) Total brood size of WT and *mafr-1* (KO) hermaphrodites (n > 10). (G) Time course showing reproductive output of WT and *mafr-1* (KO) hermaphrodites throughout reproductive span. Unless otherwise specified, statistical comparisons made by Student’s t-test. ns = no significance, * p < 0.05, ** p < 0.01, *** p < 0.001, **** p < 0.0001.

### MAFR-1 interacts with sperm-enriched proteins

MAF1 has been shown to physically interact with components of the RNA pol III complex (Desai et al., 2005; Lee et al., 2015; Oficjalska-Pham et al., 2006; Reina et al., 2006), but other direct interactors remain elusive. In order to look for novel protein interactors with MAFR-1, we performed a yeast two-hybrid screen using MAFR-1 as bait. We identified 62 putative protein-protein interactors of MAFR-1 (Table S1), which represent GO-terms comprising multiple essential cellular processes (Fig. S2*A*). Among these hits, we defined a novel class of germline- or spermatid-enriched (Yook et al.) putative MAFR-1 interactors: SSS-1, F48C1.6, and MSP-53. Because a role for MAF1 in germ cells has not been previously described, we chose these putative interactors for further analysis. We first examined the RNA expression levels of each putative interactor in wild type and *mafr-1* (KO) animals by qPCR, which revealed increased expression of *msp-53* and *sss-1*, but not *F48C1.6* relative to WT (Fig S2*B*). Next, we confirmed the physical interaction of MAFR-1 with SSS-1 (**Fig. 2*A***), and F48C1.6 (**Fig. 2*B***) biochemically by co-expression in *E.coli* followed by affinity purification MAFR-1, which facilitated the co-purification of each interactor (**Fig. 2*A*-*B***). While we observed clear enrichment for SSS-1 and F48C1.6, our ability to measure enrichment of MSP-53 was confounded by its association with the Ni-NTA resin (Fig. S2*C*); a quality likely associated with its known capacity to mediate multiple interactions (Smith, 2014; Smith and Ward, 1998). Based on their enriched expression in spermaids, we next assessed the impact of reducing the expression of SSS-1 or F48C1.6 by RNAi on sperm quality. Sperm size is a well-established quality that influence male reproductive success through sperm competition (LaMunyon and Ward, 1998). *C. elegans* males produce sperm that are significantly larger than hermaphrodite sperm and this size difference facilitates male competitive advantage when mating occurs. As expected based on their enrichment in germ cells, RNAi of each of these genes resulted in decreased spermatid size (**Fig. 2*C-D***). Unsurprisingly, reducing expression of MSP-53 through RNAi also resulted in reduced spermatid size (Fig. S2*D*). In light of these findings, we hypothesized that MAFR-1 might also function in the male germline to influence reproductive success.

**Figure 2.**
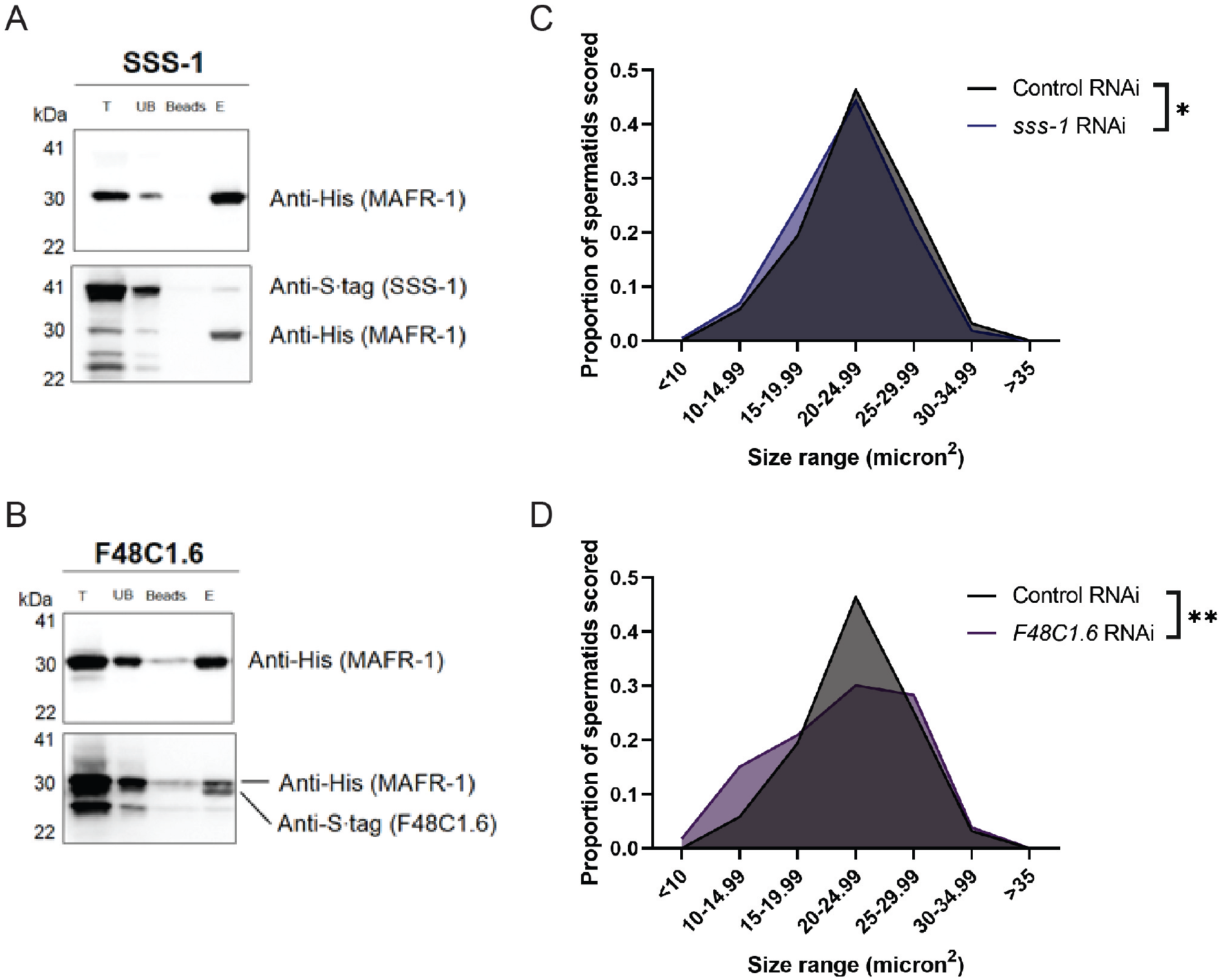
MAFR-1 interacts with SSS-1 and F48C1.6, which affect spermatid size. (A-B) Western Blot analysis of Ni-NTA column co-purification of MAFR-1 and SSS-1 (A), and MAFR-1 and F48C1.6 (B). T = total lysate, UB = unbound fraction, Beads = beads after elution, E = eluted fraction. (C-D) Spermatid size in day 1 adult males following RNAi depletion of *sss-1* (C), and *F48C1.6* (D). Experiments done in biological triplicate, statistical comparisons made by Student’s t-test. ns = no significance, * p < 0.05, ** p < 0.01, *** p < 0.001, **** p < 0.0001.

### MAFR-1 regulates male sperm fitness

To investigate the role of MAFR-1 in spermatogenesis, we looked at two metrics of spermatid quality: size and activation. We found that *mafr-1* (KO) males produced significantly smaller spermatids (**Fig. 3*A***). Next, we sought to determine the causal relationship between MAFR-1 negative regulation of RNA pol III activity and spermatid size. BRF-1 is a transcription factor required for proper RNA pol III (Buratowski and Zhou, 1992; Colbert and Hahn, 1992; Kassavetis et al., 1992; Larminie and White, 1998; Lopez-De-Leon et al., 1992). Because *mafr-1* (KO) animals have increased RNA pol III activity, we used RNAi toward *brf-1* and found that male spermatid size was restored in *mafr-1* (KO) animals (**Fig. *3B***). These results suggest a role for MAFR-1 in regulating sperm quality and competitive ability.

**Figure 3.**
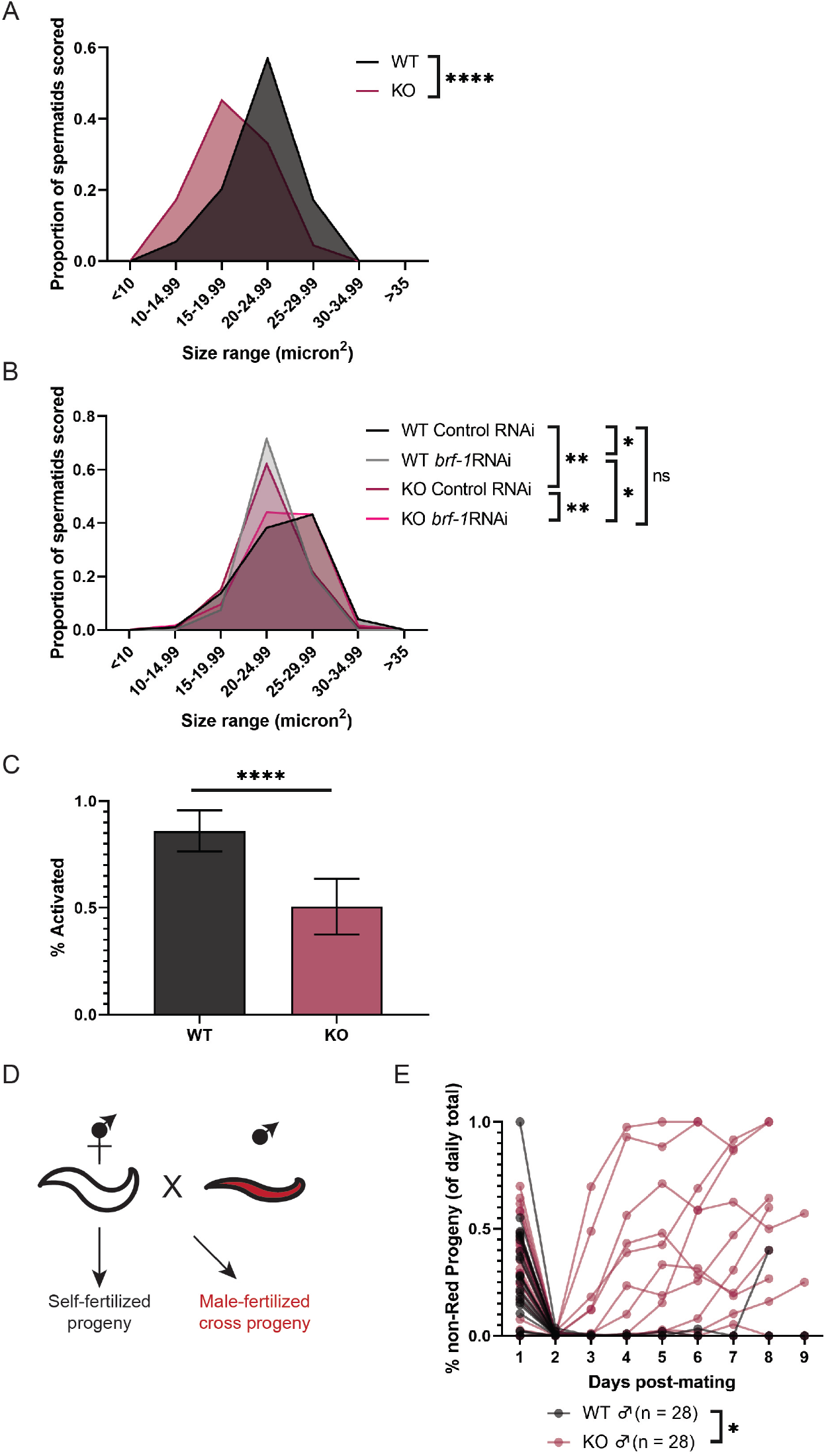
*mafr-1* (KO) males have diminished sperm quality. **(A)** Spermatid size in WT and *mafr-1* (KO) males. (B) Spermatid size of day 2 adult WT and *mafr-1* (KO) males following RNAi knockdown of *brf-1*. (C) *In vitro* Pronase activation of WT and *mafr-1* (KO) male spermatids. Experiment done in biological triplicate. Statistical comparisons made by Student’s t-test. (D) Schematic diagram showing setup of male sperm competition mating assay. (E) Proportion of progeny fertilized by self-sperm from WT and *mafr-1* (KO)-mated hermaphrodites. Experiments done in biological triplicate. Statistical comparisons made by Student’s t-test. ns = no significance, * p < 0.05, ** p < 0.01, *** p < 0.001, **** p < 0.0001.

In addition to size, the speed of male sperm is greater than hermaphrodite-derived sperm, which also enhances male competitive advantage for fertilizing an oocyte (LaMunyon and Ward). To become motile, spermatids must become activated, a sophisticated process under the control of genetic and environmental factors (Smith, 2014; Ward et al., 1983; Yen et al., 2020). In the laboratory, isolated spermatids can be activated *in vitro* by exposure to Pronase (Ward et al., 1983; Yen et al., 2020). We found that *mafr-1* (KO) male sperm are less capable of activation upon treatment with Pronase (**Fig. *3C***). Thus, *mafr-1* (KO) males have smaller sperm with a reduced capacity to mature, which could impact their ability to fertilize hermaphrodite oocytes.

In order to test the physiological consequence of *mafr-1* (KO) on sperm fitness phenotypes, we designed an assay to assess sperm quality (Yen et al., 2020). When hermaphrodites are mated to WT males, nearly all resulting progeny are derived from male sperm (LaMunyon and Ward, 1995). This can be visualized if males harboring a wrmScarlet transgene are used for mating; progeny derived from male sperm express wrmScarlet while progeny stemming from hermaphrodite self-sperm would not express wrmScarlet (**Fig. *3D***). When WT males expressing a ubiquitous wrmScarlet marker are mated to WT hermaphrodites, nearly all subsequent progeny over the reproductive span express wrmScarlet (**Fig. *3E***). In contrast, when wrmScarlet-expressing *mafr-1* (KO) males were mated to WT hermaphrodites, 10 of 28 individuals produced progeny that were non-Red (as opposed to 1 of 28 in WT males), which indicates a significant impairment of these *mafr-1* (KO) male sperm to outcompete WT hermaphrodite self-sperm. Taken together, these data reveal a novel role for MAFR-1 in spermiogenesis.

### MAFR-1 impacts sperm quality cell autonomously

One possible explanation for the discrepancy of phenotypes resulting from RNAi and genetic studies, is the lack of uniformity of RNAi across different tissues (Conte et al., 2015; Fire, 2007; Kamath et al.; Shiu and Hunter; Timmons et al.). To assess whether MAFR-1 functions cell autonomously in the germline to regulate sperm quality, we restored MAFR-1 in the germline of *mafr-1*(KO) animals, driving *mafr-1* expression with the *pie-1* promoter (Merritt et al., 2008; Merritt and Seydoux, 2010; Tenenhaus et al., 1998) (Fig. S3*A*). Animals with germline-specific rescue of *mafr-1* were generally healthy but exhibited a slight developmental delay (Fig. S3*B*). Germline-specific expression of *mafr-1* in *mafr-1* (KO) males restored spermatid size to that of WT males (**Fig. 4*A***), while partially restoring spermatid activation (**Fig. 4*B***). Importantly, germline rescue of MAFR-1 also restored the competitive ability of *mafr-1*(KO) male sperm in our mating assay, with only 2 of 25 Rescue-mated WT hermaphrodites producing non-Red progeny (**Fig. 4*C***), without affecting total brood size (Fig. S3*C*). These data suggest a cell-autonomous role for MAFR-1 in the male germline and collectively our study reveals a role for MAFR-1 in multiple parameters of sperm quality and male reproductive fitness (**Fig. 4*D***).

**Figure 4.**
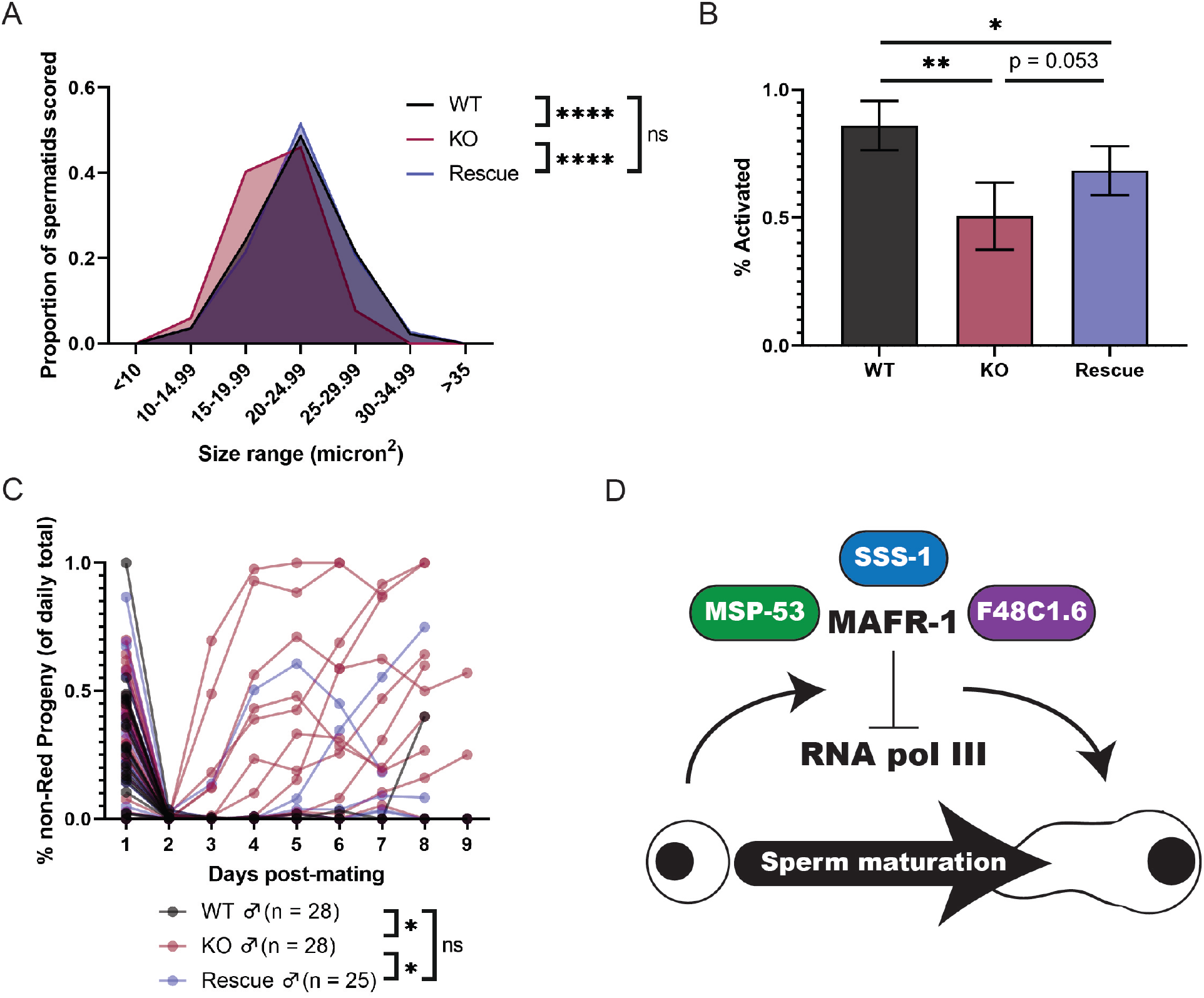
Germline expression of WT *mafr-1* rescues sperm quality phenotypes. **(A)** Spermatid size in WT, *mafr-1* (KO), and germline-specific rescue males. (B) *In vitro* Pronase activation of WT, *mafr-1* (KO), and germline-specific rescue male spermatids. *mafr-1* (KO)/Rescue comparison done via unpaired, one-tailed t-test. Experiments done in biological triplicate. (C) Proportion of progeny fertilized by self-sperm from WT, *mafr-1* (KO), and germline-specific rescue-mated hermaphrodites. (D) Model of effects of MAFR-1 on spermatid quality. MAFR-1 interacts with MSP-53, SSS-1, and F48C1.6, and its repression of RNA pol III is required for proper sperm maturation. Unless otherwise specified, statistical comparisons made by Student’s t-test. ns = no significance, * p < 0.05, ** p < 0.01, *** p < 0.001, **** p < 0.0001.

## DISCUSSION

In light of the conflicting phenotypes associated with the loss of *Maf1* by RNA interference (RNAi), genetic mutations in metazoans (Bonhoure et al., 2015; Khanna et al., 2014; Khanna et al., 2015; Pradhan et al., 2017; Rideout et al., 2012), and in cultured cell models (Palian et al., 2014), we characterized a loss-of-function allele in a second metazoan model. While *mafr-1* (KO) *C. elegans* are generally healthy (**Fig. 1*B-C***), with brood sizes comparable to WT (**Fig. 1*F***), novel interactors discovered in a yeast two-hybrid screen (**Fig. 2**, Fig. S2) prompted us to investigate the roles of MAFR-1 in the germline, specifically of males. Recent work has found MAF1 expression in embryonic stem cells drives differentiation (Chen et al., 2018). *C. elegans* possess only one population of stem cells—the germline— which originate from a single cell and become progressively more mature as they migrate through the gonad away from the distal tip cell (Kimble, 1981). In fact, previous studies have found *mafr-1* enriched in the *C. elegans* germline, but the significance of this finding has never been studied (Ebbing et al., 2018; Grun et al., 2014; Han et al., 2017).

In addition to the interaction of MAFR-1 with novel spermatid-enriched proteins, we were also prompted to examine its role in reproduction outside of vitellogenesis, as *mafr-1* (KO) animals do not replicate the vitellogenin-related phenotypes observed under *mafr-1* RNAi. While it is possible that *mafr-1* RNAi causes off-target effects that affect regulation of vitellogenin expression, RNAi effects generally require at least 25 nucleotides of high sequence similarity (Conte et al.; Parrish et al.; Rual et al.), and the longest region of sequence homology outside of *mafr-1* of the RNAi clone used is 22 nucleotides (Table S2). However, it is possible that *mafr-1* RNAi results in different regulation of vitellogenins through persistent expression, as RNAi provides significant knockdown, but not complete ablation, of gene expression. Additionally, RNAi is not uniformly effective across tissues, providing inefficient knockdown in the neurons (Kamath et al.; Timmons et al.), pharynx (Shiu and Hunter), and vulva (Conte et al.; Parrish et al.), which could be acting cell non-autonomously to cause pleiotropic gene expression phenotypes. Lastly, while the RNAi methods employed in previous work (Khanna et al.) provided efficient knockdown throughout the majority of the *C. elegans* lifespan, they were only used for a single generation, knocking down gene expression only post-embryonically. Genetic knockout results in complete removal of gene expression at all life stages, from totipotent zygote to terminally-differentiated adult cells. In the case of a transcriptional regulator, such as *mafr-1*, embryonic expression (as in single-generation RNAi) could have long-lasting developmental effects that may not be recapitulated in the genetic knockout (Chen et al.).

Prior to this study, known physical interactors of MAF1 and its homologs were limited to regulatory kinases and phosphatases (Kantidakis et al., 2010; Michels et al., 2010; Shor et al., 2010), as well as RNA pol III subunits and transcription factors (Desai et al., 2005; Lee et al., 2015; Oficjalska-Pham et al., 2006; Reina et al., 2006). Interaction of MAFR-1 with proteins expressed in a tissue-specific manner implies that, through these unique interactors, MAFR-1 may be affecting different processes in different tissues. Furthermore, restoration of WT MAFR-1 expression in the germline rescued, or partially rescued, all measured metrics of sperm quality (**Fig. 4*A-C***), indicating the cell-autonomous nature of the role of MAFR-1 in sperm maturation. To the best of our knowledge, this is the first instance of manipulation of any MAF1 homolog in any individual tissue. Taken together, our findings suggest that MAF1 may have unique roles in unique tissues that warrant further investigation.

While *mafr-1* (KO) animals have similar lifespans (**Fig. 1*C***) and developmental rates (**Fig. 1*B***) to WT, germline-rescued *mafr-1* (KO) animals take approximately two hours longer than WT or *mafr-1* (KO) to become gravid adults (Fig. S3*B*). MAF1 is activated in human cells in response to serum starvation (Goodfellow et al., 2008; Orioli et al., 2016), and confers starvation resistance in other organisms (McLean and Jacobs-Lorena, 2017; Upadhya et al., 2002). Expression of MAFR-1 under a non-endogenous promoter prevents it from being transcriptionally regulated by its normal cues, which could conceivably result in perceived starvation. In the germline, *C. elegans* have been shown to slow oogenesis in response to starvation (Seidel and Kimble, 2011), and thus, *pie-1* promoter-driven MAFR-1 expression may cause the observed delay in reproductive maturation of *mafr-1* (KO) animals through perceived starvation in the germline.

MAF1 is generally viewed as a repressor of growth (Li et al., 2016). Seemingly contrary to this paradigm, spermatids from males lacking *mafr-1* are smaller than those of WT males (**Fig. 3*A***). In *C. elegans*, as sperm size is correlated with competitive advantage, and therefore general sperm quality (LaMunyon and Ward, 1998), small sperm can be an indication of poor sperm quality. RNA pol III transcripts make up an significant fraction of spermatid RNA populations (Georgiadis et al., 2015; Ma et al., 2014), and while mature spermatids are transcriptionally and translationally quiescent (Johnson et al., 2007; Kierszenbaum and Tres, 1975), erroneous RNA pol III activity in germ cells likely results in lower quality, and therefore smaller, spermatids. Rescue of MAFR-1 expression in the germline rescued the diminished sperm quality of *mafr-1* (KO) males (**Fig. 4*A-C***), as did restoration of RNA pol III activity by RNAi of RNA pol III transcription factor *brf-1* (**Fig. 3*B***). Our results suggest that tight regulation of RNA pol III activity, through MAFR-1 or another mechanism, is vital for proper sperm function.

In summary, our work defines three novel interactors of MAFR-1, and a unique role in the male germline affecting sperm maturation, likely through its regulation of RNA pol III activity (**Fig. 4*D***).

## EXPERIMENTAL PROCEDURES

### *C. elegans* strains and maintenance

*C. elegans* were maintained at 20°C on 6 cm plates of Nematode Growth Medium (NGM) supplemented with streptomycin and seeded with OP50-1 *E. coli*. For RNAi experiments, NGM plates were supplemented with 5 mM IPTG and 100 ug/ml carbenicillin and seeded with HT115 *E. coli* expressing dsRNA targeting gene as specified.

The following strains were used: wild type (WT) N2 Bristol, PHX557[*mafr-1(syb557)* I], SPC489 [*mafr-1(syb557)* I; *laxSi01 – pie-1p∷mafr-1∷mafr-1 3’UTR cb-unc-119(+)* II], WBM1143 [*wbmIs67 – eft-3p∷3XFLAG∷wrmScarlet∷unc-54 3’UTR* V], SPC490 [*mafr-1(syb557)* I*; wbmIs67 – eft-3p∷3XFLAG∷wrmScarlet∷unc-54 3’UTR* V], SPC491 [*mafr-1(syb557)* I; *laxSi01 – pie-1p∷mafr-1∷mafr-1 3’UTR cb-unc-119(+)* II; *wbmIs67 - eft-3p∷3XFLAG∷wrmScarlet∷unc-54 3UTR* V], DH1033 [*sqt-1(sc103)* II; *bIs1 – vit-2∷GFP rol-6(su1006)* X], SPC492 [*mafr-1(syb557)* I; *sqt-1(sc103)* II; *bIs1 – vit-2∷GFP rol-6(su1006)* X]

### Yeast-2-Hybrid Screen

Bait and prey plasmids were generated by cloning *mafr-1* into pLexA and a *C. elegans* cDNA library into pACT2.2 (Addgene), respectively. Interaction was tested on synthetic complete agar that lacked leucine and tryptophan and was supplemented with X-α-gal. Interactors were identified by transforming Y2HGold (Clontech) with bait (pLexA-MAFR-1) and the cDNA prey library. Positive clones were grown in the absence of tryptophan and sequenced. Sequenced clones were then retested individually.

### Biochemical Co-purification

MAFR-1 and respective interactors were cloned into MCS1 and MCS2 of pCOLA2, respectively. Origami K-12 *E. coli* were grown to stationary phase and expression of pCOLA2 vector was induced using IPTG for two hours. Cells were pelleted from 50 ml induced culture, and frozen. Proteins were isolated and purified as in Ni-NTA Purification System protocol (Novex by Life Technologies).

### RNA extraction and gene expression

Worms were washed from plates using M9 containing 0.01% Triton-X100, washed twice in M9, and frozen at −80°C in 500 ul TRI-Reagent (Zymo). Frozen worms were thawed on ice, and cuticles manually disrupted using 25G needle. RNA was extracted using Direct-zol RNA Miniprep kit (Zymo Research R2071). cDNA was synthesized using qScript reverse transcription kit (Quanta Biosciences), diluted, and quantitative PCR was performed using PerfecTA SYBR Green (Quanta Biosciences). Primers used as previously described in Pradhan et al. and Khanna et al. (Pradhan et al., 2017). New primers used in this study are as follows:

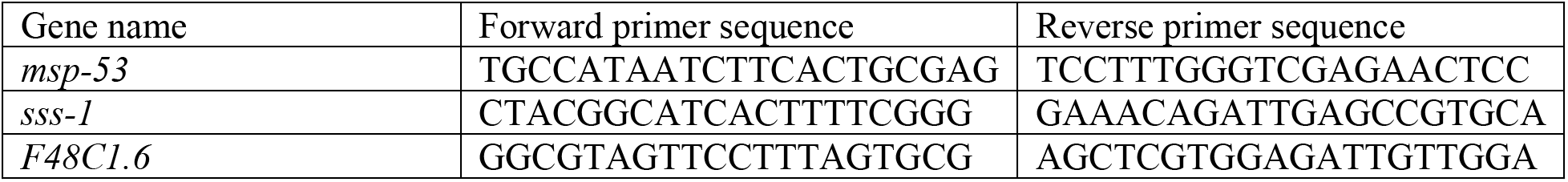

### Nile red lipid staining

Synchronous L4 animals were stained as in (Escorcia et al.). Briefly, animals were washed from plate, fixed in 40% isopropanol for 3 minutes, stained in 0.03 mg/ml Nile Red in 40% isopropanol for 2 hours, and de-stained in PBS with 0.01% Triton for 30 minutes. Stained worms were mounted in de-staining solution and imaged using DIC and GFP filters on Zeiss Axio Imager, with ZEN software. Fluorescence of individual worms was measured using ImageJ software (NIH).

### Lifespan analysis

Lifespan data was collected as previously described in the HT115 RNAi *E. coli* diet (Dalton and Curran, 2018; Haghani et al., 2019; Paek et al., 2012; Pang and Curran, 2014). In brief, plates containing synchronized populations were scored daily for survival, beginning at L4 stage. Each line represents the average of three populations of 50 animals each. Animals that died of bursting, matricide, or crawling off plate were censored.

### Reproduction

Self-reproduction: L1-synchronized hermaphrodites were grown to L4 and single worms were placed on individual plates. Worms were transferred to fresh plates every 24 hours until egg laying ceased, and progeny were counted 48 hours after hermaphrodite was removed from plate.

Mated-reproduction: Single L4 hermaphrodites were placed with virgin day 1 adult males in a 1:1 ratio on plates seeded with 20 μl OP50, and allowed to mate overnight. Males were removed from plates, and hermaphrodites were transferred to fresh plates every 24 hours until egg laying ceased. Progeny were counted and scored for wrmScarlet fluorescence 48 hours after hermaphrodite was moved from plate. Animals were censored as “not sufficiently mated” if male sperm did not suppress self-progeny production to >5% of the daily total on day 2. Hermaphrodites were counted as producing non-Red progeny if >2% of the total progeny produced after day 1 of adulthood were non-Red (progeny produced on the first day were not included due to variance in mating efficiency).

### Sperm size analysis

Five virgin day 1 adult males were dissected in SM buffer containing dextrose (Yen et al., 2020) to release spermatids. Spermatids were imaged using Zeiss Axio Imager, and size measured using ImageJ software. For experiments in which *sss-1*, *F48C1.6*, and *msp-53* RNAi were employed, animals were hatched on RNAi, and treated as OP50 animals—L4s were sequestered for 24 hours and virgin day 1 adult males were dissected.

For experiments in which *brf-1* RNAi was employed, RNAi was administered post-developmentally: animals were hatched on OP50, and L4s were treated with RNAi and sequestered for 48 hours, then virgin day 2 adult males were dissected.

### Spermatid activation assay

Five males were dissected in SM buffer containing BSA (Yen et al., 2020). An equal volume of SM buffer containing 400 μg/ml Pronase (Millipore Sigma) was added, and spermatids were allowed to activate for 15 minutes.

### VIT-2∷GFP imaging/quantification

Synchronous animals were washed from plates and fixed in 40% isopropanol for 2 hours before a 30-minute wash in PBS with 0.01% Triton-X100. Fixed animals were mounted and imaged on a Zeiss Axio Imager, and fluorescence of most proximal oocyte was quantified using ImageJ software (NIH). In order to ensure animals of the same developmental stage were quantified, only animals with exactly two embryos in utero were imaged. Fluorescence intensity per animal is equal to the sum of the corrected total cell fluorescence (CTCF) of both proximal oocytes.

### Developmental timing assay

Synchronous populations of L1 animals were dropped OP50, and beginning at 54 hours post-drop scored for gravidity. Animals with uteruses containing at least one egg were considered to be gravid.

## DATA AVAILABILITY

All data are contained within the manuscript.

## ACKNOWLEDGEMENTS

We thank J. Gonzalez and L. Thomas for technical assistance; C-A. Yen for critical reading of the manuscript. Some strains were provided by the CGC, which is funded by the NIH Office of Research Infrastructure Programs (P40 OD010440).

## AUTHOR CONTRIBUTIONS

S.P.C. designed the study; A.M.H. performed the experiments; A.M.H. and S.P.C. analyzed data. A.M.H. and S.P.C. wrote and revised the manuscript.

## FUNDING AND ADDITIONAL INFORMATION

This work was funded by the NIH R01GM109028 and R01AG058610 to S.P.C. and T32AG000037 to A.MH..

The content is solely the responsibility of the authors and does not necessarily represent the official views of the National Institutes of Health

## SUPPORTING INFORMATION

**Supplemental Figure 1.**
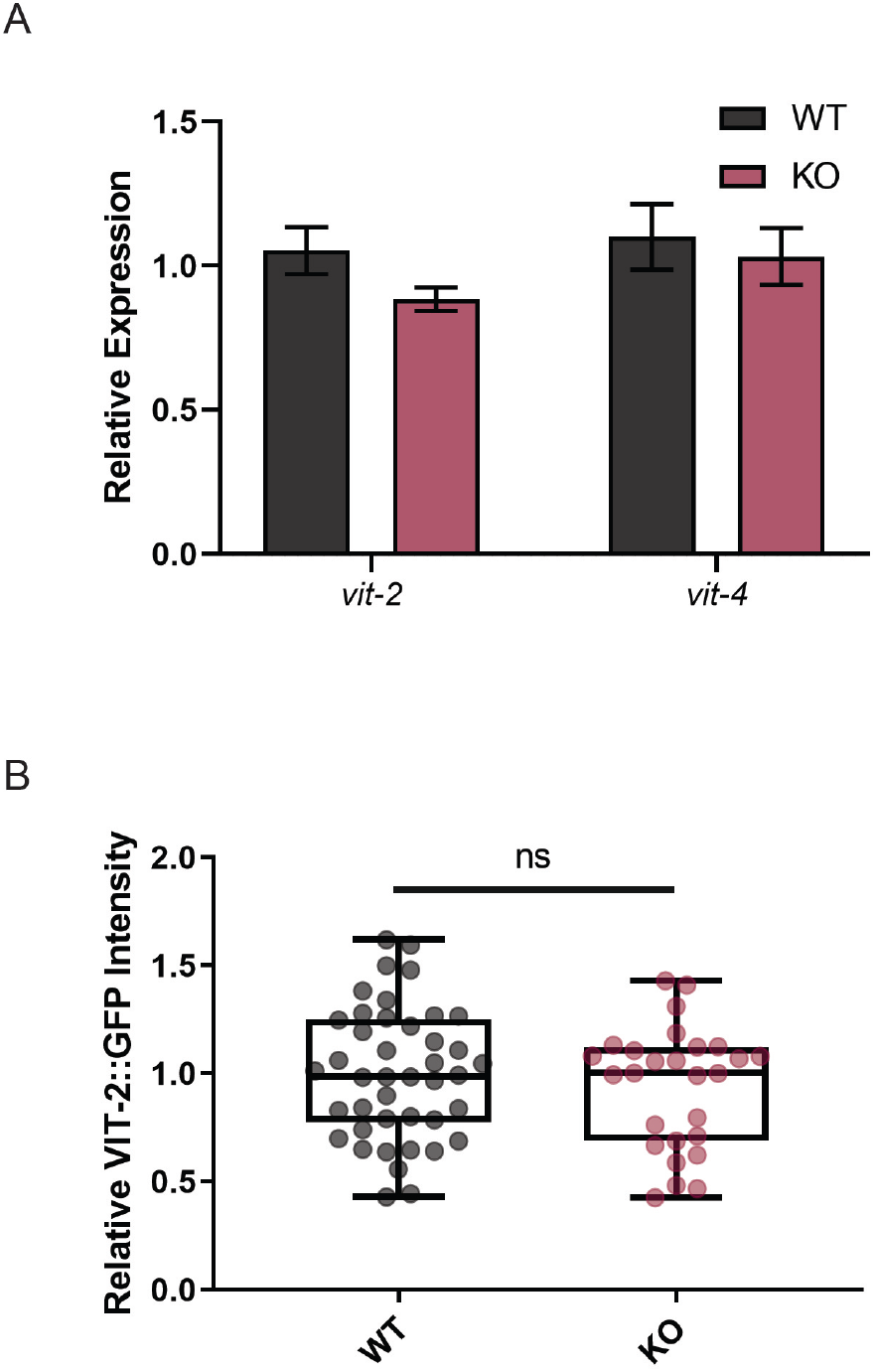
Vitellogenin expression is unaffected by *mafr-1* (KO). (A) Quantitative PCR analysis of mRNA expression of *vit-2* and *vit-4* in WT and *mafr-1* (KO) animals. No statistical difference found by Student’s t-test. (B) Quantification of VIT-2 expression by fluorescence intensity of VIT-2∷GFP fusion protein. No statistical difference found by Student’s t-test.

**Supplemental Figure 2.**
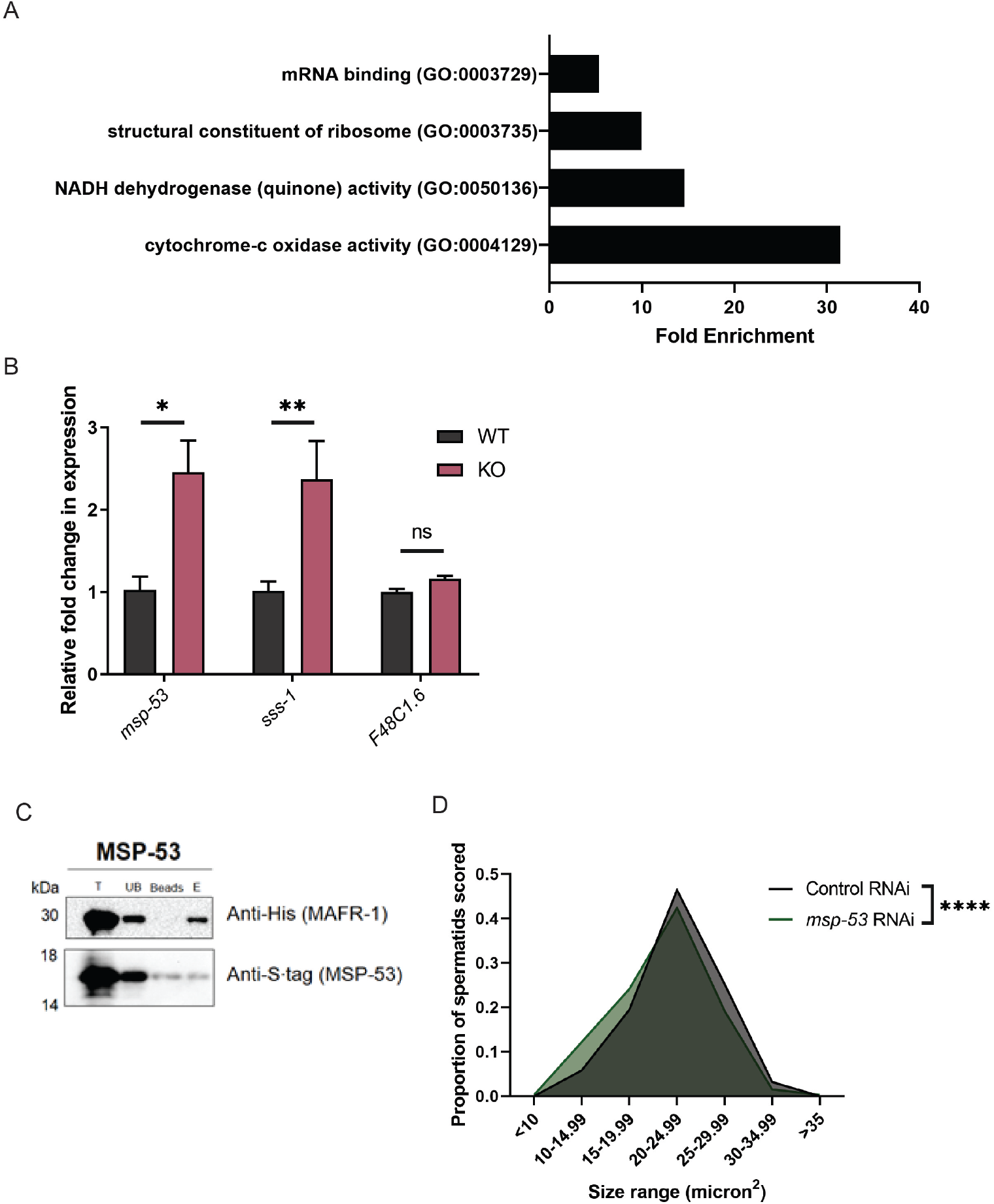
Yeast two-hybrid identified diverse classes of putative interactors, including MSP-53. (A) Molecular function GO-term enrichment analysis of Y2H positive hits. (B) Quantitative PCR expression analysis of *msp-53*, *sss-1*, and *F48C1.6* in WT and *mafr-1* (KO) animals. (C) Western Blot analysis of Ni-NTA column co-purification of MAFR-1 and MSP-53. (D) Spermatid size in day 1 adult males following RNAi depletion of *msp-53*. Experiment done in biological triplicate. Statistical comparisons made by Student’s t-test. ns = no significance, * p < 0.05, ** p < 0.01, *** p < 0.001, **** p < 0.0001.

**Supplemental Figure 3.**
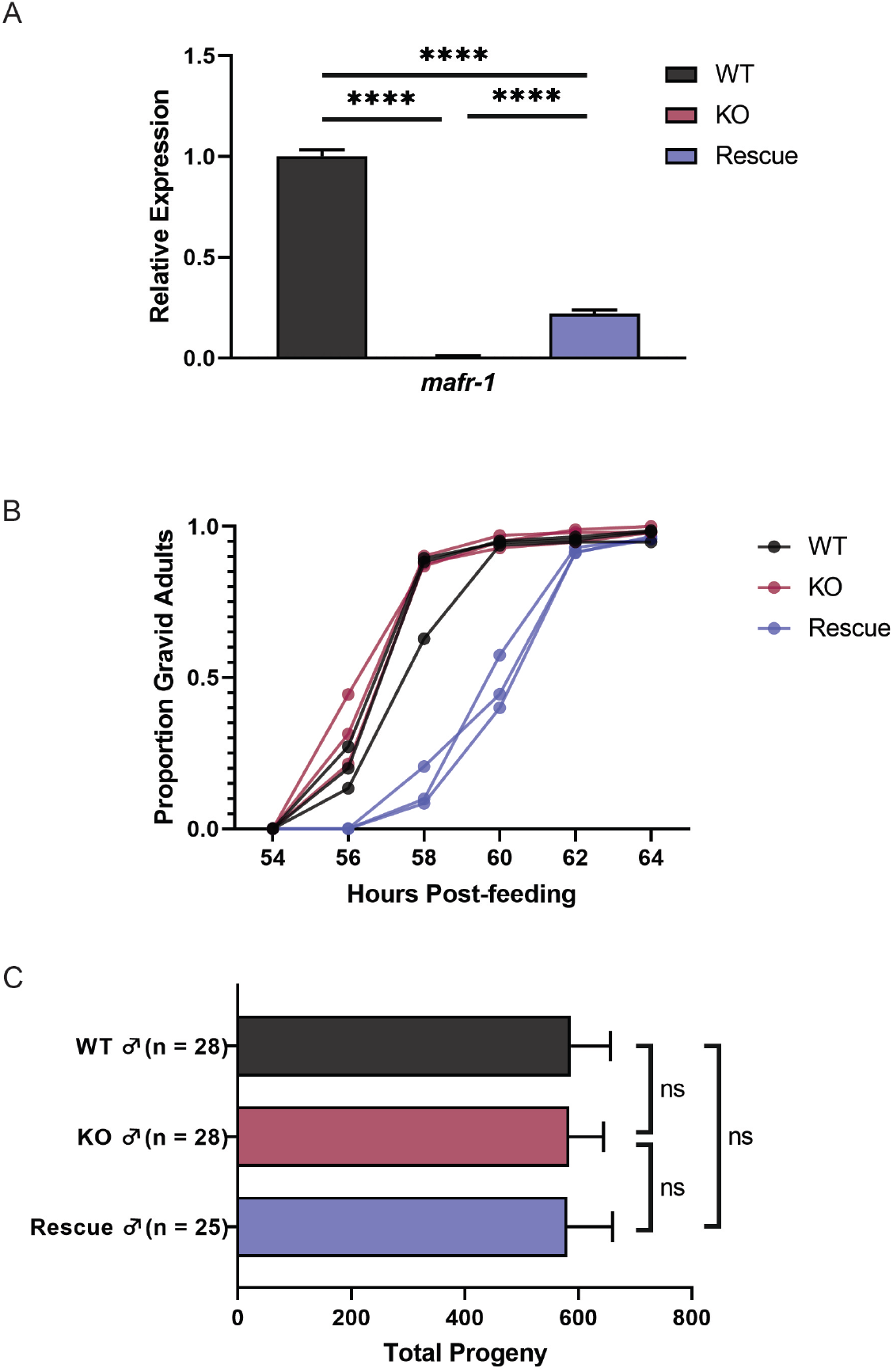
Germline-specific rescue of *mafr-1* (KO) results in developmentally slow, but fertile animals. (A) Quantitative PCR analysis of *mafr-1* expression in WT, *mafr-1* (KO), and germline-rescued animals. (B) Developmental time course of WT, *mafr-1* (KO), and germline-specific rescue hermaphrodites. (C) Total brood size of WT, *mafr-1* (KO), and germline-specific rescue-mated WT hermaphrodites. Statistical comparisons made by Student’s t-test. ns = no significance, * p < 0.05, ** p < 0.01, *** p < 0.001, **** p < 0.0001.

**Supplemental Table 1.** List of positive hits identified in yeast two-hybrid screen.

| Sequence ID | Gene Identity | # Hits |
| --- | --- | --- |
| K07E3.4 | *C. elegans* uncharacterized protein K07E3.4 | 1 |
| D1054.11 | *C. elegans* ULE-3 | 1 |
| F25B5.5 | *C. elegans* CDK5RAP1-like protein (F25B5.5) | 1 |
| C47B2.3 | *C. elegans* TBA-2 | 1 |
| C03C10.3 | *C. elegans* RNR-2 | 1 |
| T03F1.10 | *C. elegans* CLEC-53 | 1 |
| C27A2.6 | *C. elegans* DSH-2 | 1 |
| K08H10.7 | *C. elegans* RDE-1 | 1 |
| C34F6.2 | *C. elegans* COL-178 | 1 |
| K02D7.3 | *C. elegans* COL-101 | 1 |
| F54C1.7 | *C. elegans* PAT-10 | 1 |
| ZK721.2 | *C. elegans* UNC-27 | 1 |
| cTel55X.1 | *C. elegans* uncharacterized protein cTel55X.1 | 1 |
| F32B6.6 | *C. elegans* MSP-77 | 1 |
| R13H9.4 | *C. elegans* MSP-53/57 | 1 |
| F32B6.5 | *C. elegans* SSS-1 | 1 |
| Y71H2AM.20 | *C. elegans* serine/threonine-protein phosphatase 2A activator Y71H2AM.20 | 1 |
| Y71F9AM.6 | *C. elegans* TRAP-1 | 1 |
| F31E3.5 | *C. elegans* EEF-1A.1 | 1 |
| F57H12.1 | *C. elegans* ARF-3 | 1 |
| C30E10.4 | *C. elegans* GLY-20 | 1 |
| F43D9.6 | *C. elegans* URM-1 | 1 |
| T03E6.7 | *C. elegans* CPL-1 | 1 |
| ZK945.2 | *C. elegans* PAS-7 | 1 |
| T26A5.7 | *C. elegans* SET-1 | 1 |
| F45E4.2 | *C. elegans* PLP-1 | 1 |
| Y48B6A.14 | *C. elegans* HMG-1.1 | 1 |
| MTCE.4 | *C. elegans* NDFL-4 | 1 |
| E04A4.7 | *C. elegans* CYC-2.1 | 1 |
| K07A12.3 | *C. elegans* ASG-1 | 1 |
| R10E11.8 | *C. elegans* VHA-1 | 1 |
| F46F11.5 | *C. elegans* CHA-10 | 1 |
| T20D3.5 | *C. elegans* uncharacterized protein T20D3.5 | 1 |
| C14B9.10 | *C. elegans* uncharacterized protein C14B9.10 | 1 |
| F53B6.4 | *C. elegans* uncharacterized protein F53B6.4 | 1 |
| T25B9.3 | *C. elegans* uncharacterized protein T25B9.3 | 1 |
| K03D3.5 | *C. elegans* uncharacterized protein K03D3.5 | 1 |
| R10E9.2 | *C. elegans* uncharacterized protein R10E9.2 | 1 |
| D1045.10 | *C. elegans* uncharacterized protein D1054.10 | 1 |
| F48C1.6 | *C. elegans* uncharacterized protein F48C1.6 | 1 |
| T24B8.2 | *C. elegans* uncharacterized protein T24B8.2 | 1 |
| C50D2.5 | *C. elegans* uncharacterized protein C50D2.5 | 1 |
| Y62H9A.2 | *C. elegans* uncharacterized protein Y62H9A.2 | 1 |
| T01C3.3 | *C. elegans* uncharacterized protein T01C3.3 | 1 |
| M28.10 | *C. elegans* uncharacterized protein M28.10 | 1 |
| F10B5.1 | *C. elegans* RPL-10 | 2 |
| F07D10.1 | *C. elegans* RPL-11.2 | 1 |
| C32E8.2 | *C. elegans* RPL-13 | 1 |
| M01F1.2 | *C. elegans* RPL-16 | 1 |
| E04A4.8 | *C. elegans* RPL-20 | 1 |
| C27A2.2 | *C. elegans* RPL-22 | 1 |
| C53H9.1 | *C. elegans* RPL-27 | 1 |
| R11D1.8 | *C. elegans* RPL-28 | 1 |
| F37C12.4 | *C. elegans* RPL-36 | 2 |
| B0041.4 | *C. elegans* RPL-4 | 1 |
| F54C9.5 | *C. elegans* RPL-5 | 1 |
| F56F3.5 | *C. elegans* RPS-1 | 1 |
| T01C3.6 | *C. elegans* RPS-16 | 1 |
| F53A3.3 | *C. elegans* RPS-22 | 1 |
| F28D1.7 | *C. elegans* RPS-23 | 1 |
| T05E11.1 | *C. elegans* RPS-5 | 1 |
| Y37E3.8 | *C. elegans* uncharacterized protein Y37E3.8 | 1 |

**Supplemental Table 2.** Sequences with homology to *mafr-1* RNAi clone.

| Sequence ID | Gene name | Region of gene | Length | % identity |
| --- | --- | --- | --- | --- |
| F46C3.1 | *pek-1* | Exon | 19 | 19/19 (100%) |
| T06E4.21 | *T06E4.21* | Intergenic | 22 | 22/23 (95%) |
| F17E9.1/D2096.13 | *col-116/D2096.13* | Intergenic | 19 | 19/19 (100%) |
| Y48E1B.7 | *lin-38* | Intron | 22 | 22/23 (95%) |

